## Supplementary material for "Designing an irreversible metabolic switch for scalable induction of microbial chemical production": All Supplementary Notes

#### S1 Dual-transcriptional regulators controlling nutrient uptake in *E. coli*

Table 1: **Table of dual-transcriptional regulators controlling nutrient uptake in *E. coli*.** Examples are taken from EcoCyc [1]. The chemical interaction network of these systems are the same, and is illustrated in Fig. 1A.

| TF | Name | Autoregulation | Inhibited gene of uptake enzyme | Inducing nutrient |
|---|---|---|---|---|
| BetI | Betaine inhibitor | Negative | bet | Choline |
| FadR | Fatty acid degradation regulon | Negative | fadD | Oleic acid (long chain FAs) |
| GalR | Galactose repressor | Positive (repressed by complex) | galP and mglBAC operon | D-galactopyranose |
| GlcC | GlcC-glycolate DNA-binding regulator | Positive (repressed by complex) | glcA from glcDEFGBA operon | Glycolate |
| LldR | Lactate regulator | Negative and positive | lldP from lldPRD operon | S-lactate |
| NagC | N-acetylglucosamine regulator | Negative | chbBCARFG operon | N,N'-diacetylchitobiose |
| NanR | N-acetyl-neuraminic acid regulator | Constitutive | nanT from nanATEK-yhcH operon | N-acetylneuraminate |
| RbsR | Ribose repressor | Negative | rbsACB from rbsDACBKR operon | D-ribose |
| RutR | Pyrimidine utilization, rut repressor | Negative | rutG from rutABCDEFG operon | Thymine |
| TyrR | Tyrosine repressor | Negative | tyrP symporter | L-tyrosine |

#### S2 Model of fatty acid uptake and its control on growth and production

Here we describe our model of the regulation of fatty acid uptake in *E. coli*, and its application to switch from growth to production for inductions of oleic acid. We list the key assumptions underlying the formulation of our model equations, describe the biological interpretation of each model parameter, and define their respective value and source.

##### S2.1 Assumptions

i) ***Gene fadE is knocked out.*** *Why?* This means less consumption of acyl-CoA, and as shown in the study of Hartline *et al.* [2] *E. coli* DH1Δ*fadE* strain reverts back from the induced to uninduced state more slowly than its parent strain DH1. ***Implication?*** Since our aim is to develop a control system that cannot revert back to the uninduced state, studying the regulation of fatty acid uptake in DH1Δ*fadE* strain is a good starting point.

ii) ***FadD expression is only controlled by FadR.*** *Why?* In its native host, FadD expression is co-regulated by FadR and the complex CRP-cAMP (EcoCyc), during mid-exponential growth, in aerobic conditions (our growth condition of interest). FadD expression is activated by CRP-cAMP levels, but inhibited by FadR. The problem is that a common carbon source for many products of interest is glucose, and in its presence CRP-cAMP level, and so FadD expression, is very low. To make FadD expression independent of the media we therefore assume that the *fadD* promoter is replaced with a synthetic promoter that has an RNAP binidng site without the need for CRP-cAMP and an operator site for FadR inhibition. ***Implication?*** Engineering the expression of FadD in this way widens its application.

iii) ***The production phenotype is defined by the concentration of controller FadR.*** *Why?* Expression of the product synthesis enzyme(s) and synthesis of the product itself is dependent on controller FadR, and so we define the production phenotype by the concentration of FadR only. ***Implication?*** We can assess the system behaviour independent of the product of interest.

iv) ***Growth rate is linearly dependent on growth-associated enzyme $E_g$.*** *Why?* The study of Usui *et al* [3] reported that for decreases in the expression of glucose-6-phosphate isomerase (Pgi) in upper glycolysis, cell growth rate was decreased in an approximately linear fashion (Supplementary Fig. 1(a)). ***Implication?*** We capture this phenomenon and use a linear model of growth rate as a function of $E_g$:

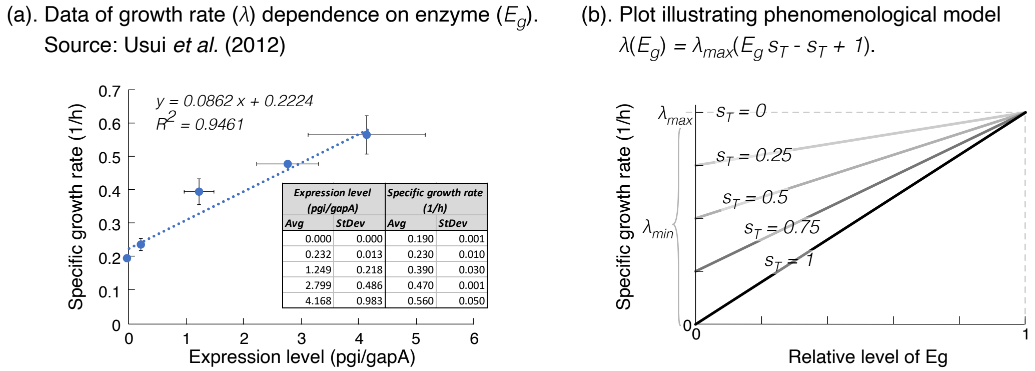

| Expression level (pgi/gapA) | | Specific growth rate (1/h) | |
|---|---|---|---|
| Avg | StDev | Avg | StDev |
| 0.000 | 0.000 | 0.190 | 0.001 |
| 0.232 | 0.013 | 0.230 | 0.010 |
| 1.249 | 0.218 | 0.390 | 0.030 |
| 2.799 | 0.486 | 0.470 | 0.001 |
| 4.168 | 0.983 | 0.560 | 0.050 |

Figure 1: **Data-driven model of growth dependence on enzyme level.** (a). Plot of the data taken form Usui *et al.* [3] showing the relation between the measured specific growth rate and concentration of a key metabolic enzyme such as glucose-6-phosphate isomerase, Pgi. Inset table of the raw values found. (b). We assume the growth-enzyme relation can be phenomenologically modelled by a linear equation, and plot this model for varying values of the parameter $s_T$ representing the severity of growth attenuation or proportional drop in growth rate.

$$\lambda(E_g) = \lambda_{\max} \cdot (E_g s_T - s_T + 1)\,, \tag{1}$$

where $s_T = 1 - \frac{\lambda_{\min}}{\lambda_{\max}}$ represents the *severity* of drop in growth rate when $E_g$ is no longer expressed, and $\lambda_{\min}$, $\lambda_{\max}$ represent growth rates at zero and maximum expression of $E_g$ (Supplementary Fig. 1(b)).

v) ***We can model enzyme expression rates in the form of a Hill function.*** *Why?* To reduce the number of free parameters in our model we assume that translation is the rate limiting step of gene expression and that ribosome-mRNA complex is at quasi-steady state. It can be shown that expression of the gene product then takes the form of a Hill function [4], either as $\frac{a\cdot(K\cdot T)^n}{1+(K\cdot T)^n}$ for $T$-activated expression or $\frac{a}{1+(K\cdot T)^n}$ for $T$-inhibited expression, where $a$ is the maximum expression rate, $K$ is the affinity of TF $T$ to bind to operator and control expression, and $n$ is the Hill coefficient. ***Implication?*** We then have a reduced model that captures the important dynamics of the system.

vi) ***No active degradation of enzymes or transcription factors.*** *Why?* As far as we are aware, the proteins in the system we study are not actively degraded and stable. Within the time length of cell doubling, dilution of proteins by cell growth and division is the primary means by which proteins are lost from the system. ***Implication?*** We model the degradation of a protein, say $E$, as dilution by growth as $\lambda \cdot E$.

#### S2.2 Equations

Our model of the endogenous system (Fig. 1(a), endogenous system) is adopted from Hartline and Mannan *et al.* [2], which models the rate of expression of TF FadR ($r_{x,R}$) and uptake enzyme FadD ($r_{x,D}$) as Hill functions, and the reaction kinetics of FadD ($r_D$) and acyl-CoA-consuming reaction PlsB ($r_B$) as Michaelis-Menten equations, to give:

$$r_{x,R} = b_R + P_R(R), \quad r_{x,D} = b_D + \frac{a_D}{1+(K_D R)^2},$$
$$r_D = \frac{k_{cat,D}\cdot \mathrm{OA}}{K_{m,D}+\mathrm{OA}} \cdot D, \quad r_B = \frac{k_{cat,B}\cdot A}{K_{m,B}+A}\cdot B.$$

Here, the expression of FadR is modelled as a function of itself to capture either its native negative autoregulation (NAR), or positive autoregulation (PAR), as engineered in [2]:

$$\mathrm{NAR}: \; P_R(R) \; = \; \frac{a_R}{1+K_R R}, \tag{2}$$
$$\mathrm{PAR}: \; P_R(R) \; = \; \frac{a_R K_R R}{1+K_R R}. \tag{3}$$

We model the sequestration of FadR by two molecules of acyl-CoA with the following mass-action kinetics:

$$r_{\mathrm{seq}} = r_f - r_r = k_f A^2 R - k_r C. \tag{4}$$

We now extend the model of Hartline and Mannan to model its application for oleic acid-inducible control on growth and production. The expression of product synthesis enzymes and the product itself does not feedback to affect the rest of the control system. Since it is exclusively dependent on FadR, we do not model production explicitly and instead define the production phenotype by a low set concentration value of FadR $R \leq 0.0033\mu$M, which means the growth phenotype is defined as $R > 0.0033\mu$M (assumption (iii)).

We do however need to model control on growth as this feeds back to affect dilution of all intracellular species. We exploit that FadR is natively a dual transcriptional regulator [5], and model FadR-activated expression rate of a growth-associated enzyme $E_g$ as a Hill function (assumption (v)):

$$r_{x,E_g} = \frac{a_g K_g R}{1+K_g R}.$$

To model how growth rate is in turn affected by $E_g$ we used experimental data from Usui, *et al.* [3] to derive a linear model that captures the linear dependence of growth rate on $E_g$ (Supplementary Equation (1), assumption (iv)).

We now write down the system of ordinary differential equations that model the rate of change of FadR

($R$), FadD ($D$), acyl-CoA ($A$), sequestered complex ($C$) and growth-associated enzyme ($E_g$):

$$\begin{aligned}
\frac{dR}{dt} &= r_{x,R} - r_{\text{seq}} - \lambda(E_g)R, \\
\frac{dD}{dt} &= r_{x,D} - \lambda(E_g)D, \\
\frac{dA}{dt} &= r_D - r_B - 2\cdot(r_{\text{seq}}) - \lambda(E_g)A, \\
\frac{dC}{dt} &= r_{\text{seq}} - \lambda(E_g)C, \\
\frac{dE_g}{dt} &= \frac{a_g K_g R}{1+K_g R} - \lambda(E_g)\cdot E_g,
\end{aligned} \tag{5}$$

where $\lambda(E_g)$ is given in Supplementary Equation (1).

##### S2.3 Parameters

The model parameters are taken from the Hartline and Mannan *et al.* model [2], but a full description of parameters, their values, and underlying assumptions explaining why we chose those values are detailed in Supplementary Table 2. It is important to note that to allow a fair comparison between the systems with NAR and PAR, we have to chose parameters so the steady state concentration of FadR prior to induction is the same in both circuits. To ensure this we scaled FadR's affinity by 7-fold to activate its own promoter ($K_R$ in Supplementary Equation (3)) (see Supplementary Table 2).

Table 2: **Parameters of the model of the system shown in Fig. 1(a).** Description and values of the parameters of our model of the control system illustrated in Fig. 1(a). The values are based on characterised parts from the literature, with reference to the source. † indicates parameters of FadR expression when under PAR.

| **Parameter description** | **Term** | **Value** | **Units** | **Reference** |
|---|---|---|---|---|
| Growth rate | $\lambda_{\max}$ | 0.1818 | $h^{-1}$ | [2], *E. coli* DH1Δ*fadE* strain. |
| Proportional drop in growth | $s_T$ | 0.7 | - | Determined from [3], *pgi* down-regulation. |
| FadR leaky expression (NAR) | $b_R$ | 0.0007 | $\mu M\cdot h^{-1}$ | [2] |
| FadR promoter strength (NAR) | $a_R$ | 0.0131 | $\mu M\cdot h^{-1}$ | [2] |
| FadR affinity to own promoter (NAR) | $K_R$ | 4.3222 | $\mu M^{-1}$ | [2] |
| FadR leaky expression (PAR) | $b_R$ | 0.0007 † | $\mu M\cdot h^{-1}$ | Assumed same as params for FadR with NAR. |
| FadR promoter strength (PAR) | $a_R$ | 0.0131 † | $\mu M\cdot h^{-1}$ | Assumed same as params for FadR with NAR. |
| FadR affinity to own promoter (PAR) | $K_R$ | 4.3222 † × 7 | $\mu M^{-1}$ | Assumed same as for FadR with NAR, except scaled so steady state FadR approx same as system with NAR before induction (OA = 0$\mu$M). |
| FadD leaky expression | $b_D$ | 0.0108 | $\mu M\cdot h^{-1}$ | [2] |
| FadD promoter strength | $a_D$ | 0.0517 | $\mu M\cdot h^{-1}$ | [2] |
| FadR affinity to *fadD* | $K_D$ | 305.95 | $\mu M^{-1}$ | [2] |
| FadD turnover rate | $k_{\text{cat,D}}$ | 49 | $h^{-1}$ | [2] |
| Michaelis constant | $K_{\text{m,D}}$ | 0.0672 | $\mu M$ | [2] |
| PlsB turnover rate | $k_{\text{cat,B}}$ | 192.91 | $h^{-1}$ | [2] |
| Michaelis constant | $K_{\text{m,B}}$ | 45429 | $\mu M$ | [2] |
| PlsB concentration | $B$ | 0.1369 | $\mu M$ | [2] |
| Forward sequestration | $k_f$ | 612.55 | $\mu M^{-2}\cdot h^{-1}$ | [2] |
| Reverse sequestration | $k_r$ | 900.73 | $h^{-1}$ | [2] |
| $E_g$ promoter strength | $a_g$ | $\lambda_{\max}$ | $\mu M\cdot h^{-1}$ | This study, to ensure full expression at $\lambda_{\max}$. |
| FadR affinity to $E_g$ promoter | $K_g$ | $=K_R=$ 4.9114 | $\mu M^{-1}$ | In this study †Supplementary Note S7.2, from fitting of data from [2]. Assumed designed with same promoter as *fadR* with PAR, for FadR-activated expression. |
| Affinity of FadR for prod synthesis enzyme $E_p$ | $K_p$ | $=K_D=$ 305.95 | $\mu M\cdot h^{-1}$ | Assume designed with FadR operator site from *fadD* promoter; value from [2]. |

#### S2.4 Determining the steady states and their stability

To characterise the control system behaviour we look at the dose-response, that is the system steady state achieved at different inducer concentrations. Here we describe how we determine the steady state(s) for a given inducer concentration and its stability.

To determine the steady states, we evaluate the system when all derivative become zero. Evaluating $dE_g/dt = 0$, $dC/dt = 0$ and $dD/dt = 0$ it can be shown that the steady states of $E_g$, $C$ and $D$ are:

$$\begin{aligned} E_g(R) &= \frac{\lambda_{\max}(s_T-1) + \sqrt{\lambda_{\max}^2(s_T-1)^2 + \frac{4\lambda_{\max}s_T a_g K_g R}{1+K_g R}}}{2\lambda_{\max}s_T}, && (6)\\ C(A,R) &= \frac{k_f A^2 R}{k_r \lambda(R)}, && (7)\\ D(R) &= \frac{1}{\lambda(R)} \cdot \left(b_D + \frac{a_D}{1+(K_D R)^2}\right), && (8) \end{aligned}$$

where $\lambda(R)$ is defined in Supplementary Equation (1). Evaluating $dR/dt = 0$, we derive first nullcline $N_1$:

$$N_1(A,R) : (b_R + P_R(R) - \lambda(R)R) \cdot \left(\frac{k_r + \lambda(R)}{k_f \lambda(R) R}\right) - A^2 = 0, \quad (9)$$

where $P_R(R)$ is defined in Supplementary Equation (6). Evaluating $dA/dt = 0$ derived second nullcline $N_2$:

$$N_2(A,R) : \frac{k_{cat,D} \cdot \mathrm{OA} \cdot D}{K_{m,D} + \mathrm{OA}} - \frac{k_{cat,B} \cdot A \cdot B}{K_{m,B} + A} - 2 \cdot \left(\frac{k_f A^2 R \lambda(R)}{k_r + \lambda(R)}\right) - \lambda A = 0. \quad (10)$$

The steady states of our system lie at the intersections of the two nullclines, which are functions completely in terms of FadR $R$ (noting that $A = A(R)$ from $N_1$). Therefore, the steady state of all other species can be derived from the steady state of $R$. It is difficult to derivate an analytical form of the steady state of $R$ only in terms of the parameters, and so we solve for it numerically using MATLAB R2018a.

However, in an effort to limit the search space over which to solve for $R$, we look for an expression of its maximum value. $R$ is maximum in the absence of OA, so when OA $= 0\mu$M then $A = 0\mu$M, and so $N_2 = 0$ and $N_1$ gives us $(b_R + P_R(R) - \lambda(R)R) = 0$. We solved this equation using local optimization solver `fminbnd` in MATLAB to give us an estimate for the maximum possible value of $R \in (0, 1000]$, for a given set of parameter values.

We determined the stability of steady states by looking at the signs of the eigenvalues of the Jacobian evaluated at the steady state, by solving its characteristic polynomial. To find the Jacobian of our system, we first wrote down our model substituting $\lambda$ with $\lambda(E_g) = \lambda_{\max} \cdot (E_g s_T - s_T + 1)$ from Supplementary Equation (8) into Supplementary Equation (5). We then evaluated the partial derivatives and found our Jacobian as:

$$J = \begin{bmatrix} P'_R - k_f A^2 - \lambda(E_g), & 0, & -2k_f AR, & k_r, & -\lambda_{\max} R s_T \\ -\frac{2a_D K_D^2 R}{(1+(K_D R)^2)^2}, & -\lambda(E_g), & 0, & 0, & -D\lambda_{\max} s_T \\ -2k_f A^2, & \frac{k_{\mathrm{cat,D}}\mathrm{OA}}{K_{\mathrm{m,D}}+\mathrm{OA}}, & -\frac{k_{\mathrm{cat,B}} B K_{\mathrm{m,B}}}{(K_{\mathrm{m,B}}+A)^2} - 4k_f AR - \lambda(E_g), & 2k_r, & -A\lambda_{\max} s_T \\ k_f A^2, & 0, & 2k_f AR, & -k_r - \lambda(E_g), & -C\lambda_{\max} s_T \\ \frac{a_g K_g}{(1+K_g R)^2}, & 0, & 0, & 0, & -2\lambda_{\max} E_g s_T + \lambda_{\max}(s_T - 1) \end{bmatrix}, \quad (11)$$

where $\lambda(E_g) = \lambda_{\max} \cdot (E_g s_T - s_T + 1)$ , and $P'(R)$ is:

$$P'(R) = -\frac{a_R K_R}{(1+K_R R)^2}, \text{ for NAR; and } P'(R) = \frac{a_R K_R}{(1+K_R R)^2}, \text{ for PAR.}$$

### S3 Sensitivity analysis of dose-response to model parameters

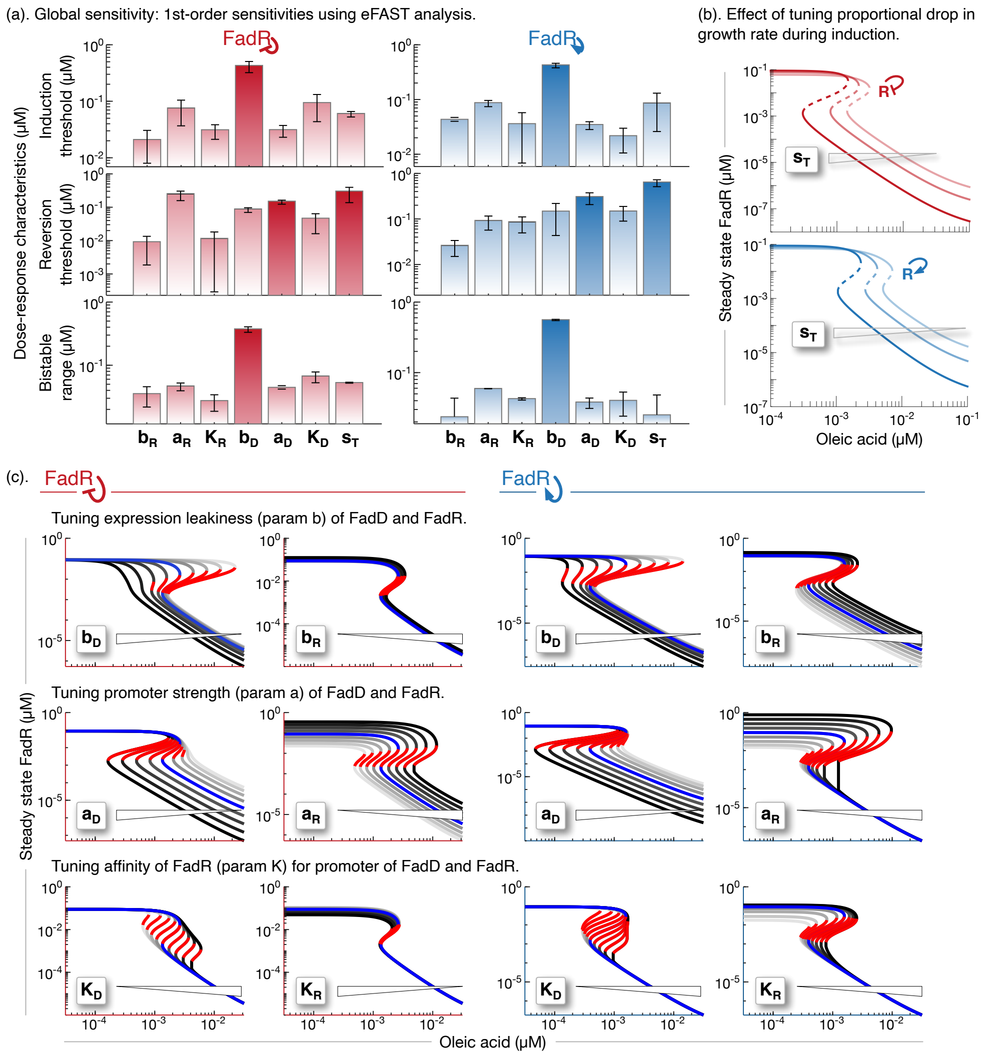

Figure 2: **Global and local sensitivity analysis of dose-response to parameter variations.** (a). Plots of average and standard deviation (error bars) of first-order sensitivity indices, based on 300 sets of parameter samples, using GSA method eFAST [6]. eFAST was implemented in MATLAB R2018a and explored values of parameters $b_R, a_R, K_R, b_D, a_D, K_D$ between 10% to 500% of their respective nominal values in Supplementary Table 2. $s_T$ was varied between 0.1–0.9. We highlight those parameters that dose-response of both circuits was most sensitive to. (b). Plots of how dose-response changed for changes in proportional drop in growth rate during induction ($s_T$), for $s_T =$ (0.23,0.49,0.75). (c). Plots of how dose-response changed for changes in one parameter at a time. We sampled nine, log-spaced values of each parameter, from 10% to 1000% of their nominal value from Supplementary Table 2. Blue curve is of nominal value.

### S4 Mathematical analysis: reversion rate depends on mode of autoregulation

Simulations of the induction dynamics showed the control systems were converging back towards the uninduced state after a temporary induction, what we refer to as reversion. We observed that reversion of the system with FadR PAR takes longer than system with FadR NAR (Fig. 3c).

***Aim:* To understand, if the slower reversion is a fundamental property of the system with PAR vs NAR, as opposed to simply a parameter choice.**

To address this aim, we looked for an analytical expression of the rate of convergence to the uninduced steady state of the two systems. We define this as the least negative eigenvalue of the Jacobian of the system evaluated at the uninduced steady state, where the Jacobian is defined in Supplementary Equation (11). We define the uninduced state as that steady state where oleic acid OA $= 0\mu$M and $(R, D, A, C, E_g) = (R, D, 0, 0, 1)$. Here, the concentrations of $R$ and $D$ are $> 0$, but those of $A$ and $C$ are $0\mu$M. Also, $E_g$ is fully expressed, and so its relative level is 1. Since the analysis of the eigenvalues is close to the uninduced steady state, we assume that there is insignificant change in growth for small enough perturbations, and so fix $s_T = 0$ and $\lambda_{\text{max}} = \widetilde{\lambda}$.

Evaluating the Jacobian $J$ in Supplementary Equation (11) at the uninduced steady state and fixed growth rate, gives:

$$J = \begin{bmatrix} P'_R - \widetilde{\lambda}, & 0, & 0, & k_r, & 0 \\ -\frac{2a_D K_D^2 R}{(1+(K_D R)^2)^2}, & -\widetilde{\lambda} & 0 & 0 & 0 \\ 0, & 0, & -\frac{k_{\text{cat,B}} B}{K_{\text{m,B}}} - \widetilde{\lambda} & 2k_r & 0 \\ 0 & 0 & 0 & -k_r - \widetilde{\lambda} & 0 \\ \frac{a_g K_g}{(1+K_g R)^2} & 0 & 0 & 0 & -\widetilde{\lambda} \end{bmatrix}.$$

Solving for the eigenvalues of the system with NAR ($e_{\text{NAR}}$) and PAR ($e_{\text{PAR}}$), we find:

$$\begin{aligned} e_{\text{NAR}} &= \left(-\widetilde{\lambda}, -\widetilde{\lambda}, -\widetilde{\lambda} - k_r, -\widetilde{\lambda} - \frac{a_R K_R}{(1+K_R R)^2}, -\widetilde{\lambda} - \frac{k_{\text{cat,B}} B}{K_{\text{m,B}}}\right)^T, \\ e_{\text{PAR}} &= \left(-\widetilde{\lambda}, -\widetilde{\lambda}, -\widetilde{\lambda} - k_r, -\widetilde{\lambda} + \frac{a_R K_R}{(1+K_R R)^2}, -\widetilde{\lambda} - \frac{k_{\text{cat,B}} B}{K_{\text{m,B}}}\right)^T. \end{aligned}$$

We find the least negative eigenvalues of the systems with NAR and PAR are:

$$\text{LNE}_{\text{N}} = -\widetilde{\lambda}; \text{ and } \text{LNE}_{\text{P}} = -\widetilde{\lambda} + \frac{a_R K_R}{(1+K_R R)^2}.$$

Since $\text{LNE}_{\text{P}} < \text{LNE}_{\text{N}}$ for non-zero parameters, then reversion rate of the system with PAR is always slower than that of the system with NAR. This phenomenon is thus a fundamental property of the circuit topology, as opposed to a choice of parameters.

#### S5 How tuning control parameters affects performance of induction dynamics

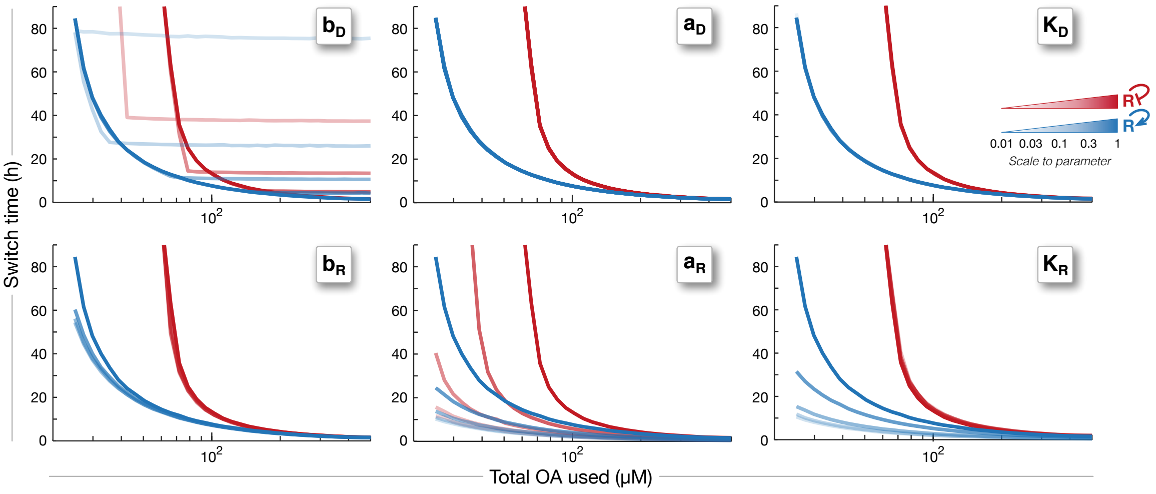

Figure 3: **Effect of varying some control parameters on induction process performance.** Plots of curves showing the performance of the induction dynamics, that is how total OA used and switch times, changed for variations in OA feed-in flux from 0.25–5$\mu$M$\cdot$ h$^{-1}$, as measured from simulations of the induction dynamics over 100 hours of the process. Different curves are generate by scaling each control circuit parameter of the circuit with NAR (red) and PAR (blue), by (0.01,0.03,0.01,0.3,1) time their respective nominal values from Supplementary Table 2 (from most faint to most opaque curves). The five different curves are difficult to see for some parameter values as they did not change much (on top of one another).

### S6 Mathematical analysis: modifying params or autoregulation only cannot enable irreversible behaviour

***Aim:*** **To determine if any of the control circuit designs considered (Supplementary Fig. 4(a)–(c)) can be engineered as an irreversible switch, that is can we find a parameter regime for this, if at all.**

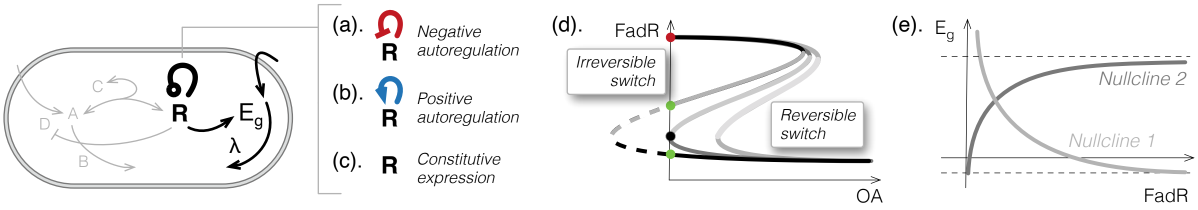

Figure 4: **Control circuits in the absence of inducer.** Schematics of the control circuit with negative autoregulation (a), positive autoregulation (b) and where FadR is constitutively expressed (c). To elucidate the steady state of the system in absence of inducing oleic acid (OA), we highlight only that part of the system that needs to be considered (black lines and text), which effectively becomes independent of the other parts (grey lines and text). (d). To determine if our system can be engineered to behave irreversibly we look at the dose-response and identify the number of steady states when OA = 0$\mu$M, i.e. cross the y-axis. If the switch is irreversible then there is at least two steady states (red and black, or red and green dots), otherwise there is only one (only red dot). (e). Sketch of the system nullclines indicates the existence of a single steady state, as proved below.

To address our aim, it is sufficient to consider the number of steady states of the system possible in the absence of oleic acid. We define an irreversible switch as that which has at least two steady states when OA = 0$\mu$M (Supplementary Fig. 4(d)). No oleic acid also means no acyl-CoA or sequestered complex too ($A = C = 0\mu$M). So, the dynamics of the system can be collapsed to considering only the dynamics of FadR ($R$) and growth associated enzyme ($E_g$), which we model as:

$$\begin{aligned} \frac{dR}{dt} &= b_R + P_R(R) - \lambda_{\max}(E_g s_T - s_T + 1)R, \\ \frac{dE_g}{dt} &= \frac{a_g K_g R}{1 + K_g R} - \lambda_{\max}(E_g s_T - s_T + 1)E_g, \end{aligned}$$

where we define $P_R(R)$ for the systems with negative autoregulation (NAR, Supplementary Fig. 4(a)), positive autoregulation (PAR), Supplementary Fig. 4(b) and constitutive expression of FadR (CON, Supplementary Fig. 4(c)) as:

$$P_{R,\text{NAR}} = \frac{a_R}{1 + K_R R}; \; P_{R,\text{PAR}} = \frac{a_R K_R R}{1 + K_R R}; \; P_{R,\text{CON}} = 0.$$

We now determine the nullclines of this reduced system and identify the number of times they intersect, that is defined as the number of steady states.

To determine the nullclines we evaluate the system at steady state. It can be shown that from an evaluation of $\frac{dR}{dt} = 0$ we derived the first nullcline $N_1$

$$N_1(R) = E_g(R) = \frac{b_R + P_R(R)}{\lambda_{\max} s_T R} - \frac{1}{s_T} + 1, \tag{12}$$

and from an evaluation of $\frac{dE_g}{dt} = 0$ we derived the second nullcline $N_2$:

$$N_2(R) = E_g(R) = \frac{\lambda_{\max}(s_T - 1) + \sqrt{\lambda_{\max}^2 (s_T - 1)^2 + \frac{4\lambda_{\max} s_T a_g K_g R}{1 + K_g R}}}{2\lambda_{\max} s_T}. \tag{13}$$

We now take a geometric approach to proving that the nullclines intercept exactly once, using Theorem 1.

**Theorem 1.** *Let $f(x)$ and $g(x)$ be continuous and differentiable functions on the interval $[a, b]$. If $f(a) < g(a)$ and $f(b) > g(b)$, and $f'(x) > 0$ and $g'(x) < 0$ (are monotonic) for $x \in [a, b]$, then $f(x)$ and $g(x)$ intercept exactly once.*

*Proof.* Let's define $h(x) = f(x) - g(x)$, so $h(x)$ is also continuous and differentiable on interval $[a, b]$. Evaluating the function at the interval ends we have:

$$\begin{aligned} h(a) &= f(a) - g(a) < 0, \\ h(b) &= f(b) - g(b) > 0. \end{aligned}$$

By the intermediate value theorem $\exists$ at least one root of $h(x) \in (a, b)$.

Let's now suppose that there exists at least two roots, and we define two of these as $c$ and $d$, so $h(c) = 0$ and $h(d) = 0$ for $\{c, d\} \in (a, b)$. By Rolle's theorem, for some point $\widetilde{c} \in (c, d)$, $h'(\widetilde{c}) = 0$.

However, $h'(x) = f'(x) - g'(x) > 0$, meaning $h(x)$ is monotonically increasing in $[a, b]$. Since $\widetilde{c} \in (c, d) \subset (a, b)$, then:
$\implies h'(\widetilde{c}) \neq 0$;
$\implies \exists$ exactly one root of $h(x) = 0$ in $(a, b) \implies f(x)$ and $g(x)$ intercept exactly once. (QED) □

To apply Theorem 1, we must first determine the properties of the two nullclines. We state that the nullclines $N_1(R)$ and $N_2(R)$, defined in the domain $R \in (0, \infty)$, are continuous and differentiable.

We first evaluate their derivatives, and after some manipulations it can be shown that:

$$\begin{aligned} \frac{dN_1}{dR} &= \begin{cases} -\frac{b_r}{\lambda_{\max} s_T R^2} & < 0, \text{ for system with CON}, \\ -\frac{b_R(1+K_R R)^2 + 2a_R K_R R + a_R}{\lambda_{\max} s_T R^2 (1=K_R R)^2} & < 0, \text{ for system with NAR}, \\ -\frac{b_R(1+K_R R)^2 + a_R K_R^2 R^2}{\lambda_{\max} s_T R^2 (1=K_R R)^2} & < 0, \text{ for system with PAR}, \end{cases} \\ \frac{dN_2}{dR} &= \frac{a_g K_g}{(1+KgR)^2 \sqrt{\lambda_{\max}^2 (s_T - 1)^2 + \frac{4\lambda_{\max} s_T a_g K_g R}{1+K_g R}}} > 0, \ \forall\, R \in (0, \infty). \end{aligned}$$

We then evaluate the nullclines at the limits of the domain, and find:

$$\begin{aligned} \lim_{R \to 0} N_1 &\to \infty, \\ \lim_{R \to 0} N_2 &= 1 - \frac{1}{s_T} \leq 0, \text{ for } s_T \in (0, 1], \\ \lim_{R \to \infty} N_1 &= 1 - \frac{1}{s_T} \leq 0, \\ \lim_{R \to \infty} N_2 &= \frac{\lambda_{\max}(s_T - 1) + \sqrt{\lambda_{\max}^2 (s_T - 1)^2 + 4\lambda_{\max} s_T a_g}}{2\lambda_{\max} s_T} > 0, \text{ for } s_T \in (0, 1]. \end{aligned}$$

So in summary, we have:

$$\begin{aligned} \lim_{R \to 0} N_2 &< \lim_{R \to 0} N_1, \\ \lim_{R \to \infty} N_2 &> \lim_{R \to \infty} N_1, \\ N_1' &< 0 \text{ irrespective of the circuit topology}, \\ N_2' &> 0. \end{aligned}$$

Then by Theorem 1, $N_1$ and $N_2$ intercept exactly once in the domain $R \in (0, \infty)$. Therefore, there is only and exactly one steady state of FadR ($R$) when OA $= 0\mu$M, and so control circuits illustrated in Supplementary Fig. 4(a), (b) and (c) cannot be engineered to behave as irreversible switches. Our sketch of the nullclines on the $(R, E_g)$-plane is given in Supplementary Fig. 4(e), and shows existence of the single steady state.

### S7 Modelling proposed control circuit and how to tune it to behave irreversibly

#### S7.1 Model formulation

We extend our mathematical model to now include the mutual inhibition between FadR and TetR (Fig. 3(a)). We make two modifications: (i) we include the dynamic expression of TetR, which we express as a Hill function of FadR, and (ii) modify the formula of the expression of FadR $P_R(R,T)$, which is now a function of both TetR and FadR. Our model is now:

$$\begin{aligned}
\frac{dR}{dt} &= b_R + P_R(R,T) - k_f A^2 R + k_r C - \lambda R, \\
\frac{dD}{dt} &= b_D + \frac{a_D}{1+(K_D R)^2} - \lambda D, \\
\frac{dA}{dt} &= \frac{k_{cat,D} \cdot \mathrm{OA}}{K_{m,D} + \mathrm{OA}} \cdot D - \frac{k_{cat,B} \cdot A}{K_{m,B} + A} \cdot B - 2 \cdot \left(k_f A^2 R - k_r C\right) - \lambda A, \\
\frac{dC}{dt} &= k_f A^2 R - k_r C - \lambda C, \\
\frac{dE_g}{dt} &= \frac{a_g K_g R}{1 + K_g R} - \lambda E_g, \\
\frac{dT}{dt} &= b_T + \frac{a_T}{1+(K_{Ri} R)^2} - \lambda T,
\end{aligned} \tag{14}$$

where growth rate $\lambda$ is given by Supplementary Equation (1), and the final ODE in Supplementary Equation (14) models the dynamics of TetR expression.

For the second modification to our model, we consider how FadR expression is controlled, and derive the subsequent formula for $P_R(R,T)$. There are three possible ways in which FadR expression is regulated by itself and TetR: (i) where there is no FadR PAR and FadR expression is only inhibited by TetR (NoPAR); (ii) where the operators for FadR binding to active its expression and TetR binding to inhibit its expression are arranged so their binding is competitive (Comp) or (iii) their binding is non-competitive (NonComp) (Fig. 3(a)).

For configuration NoPAR, we assume that TetR binds to two *tetO* operator sites, one between the -35 and -10 RNAP binding sites and the other towards the start codon, as arranged in the $P_A$ promoter from [7]. Writing down the dynamics of DNA availability, binding of TetR to operators, mRNA synthesis, and FadR synthesis as mass-action kinetics, but then assuming constant total DNA and rapid binding-unbinding of DNA and TetR, i.e. quasi-steady state of the bound complex, we can derive the FadR synthesis rate from first principles, which can be shown to be:

$$\text{NoPAR}: \; P_R(T) = \frac{a_R}{1+(K_T T)^2}. \tag{15}$$

For the circuitry where both FadR and TetR control the expression of FadR but bind competitively (Comp), we can write down the dynamics of free DNA (DNA), mRNA ($m$), the DNA complexes with TetR ($c_{d,t}$) and FadR ($c_{d,r}$), and FadR ($R$) with mass action kinetics as follows:

$$\begin{aligned}
\frac{d\mathrm{DNA}}{dt} &= -k_{f,t}^2 \cdot T^2 \cdot \mathrm{DNA} - k_{f,r} \cdot R \cdot \mathrm{DNA} + k_{r,t} \cdot c_{d,t} + k_{r,r} \cdot c_{d,r} \\
\frac{dm}{dt} &= s_m \cdot c_{d,r} - \delta_m \cdot m \\
\frac{dc_{d,t}}{dt} &= k_{f,t}^2 \cdot T^2 \cdot \mathrm{DNA} - k_{r,t} \cdot c_{d,t} \\
\frac{dc_{d,r}}{dt} &= k_{f,r} \cdot R \cdot \mathrm{DNA} - k_{r,r} \cdot c_{d,r} \\
\frac{dR}{dt} &= s_r \cdot m - \delta_r \cdot R.
\end{aligned} \tag{16}$$

Similarly, for the circuitry where FadR and TetR binding non-competitively (NonComp), the dynamics of free DNA (DNA), mRNA ($m$), the DNA complexes with TetR ($c_{d,t}$), FadR ($c_{d,r}$), and now also both ($c_{d,t,r}$), and FadR ($R$) can be written down with mass action kinetics as follows:

$$\begin{aligned}
\frac{d\text{DNA}}{dt} &= -k_{f,t}^2 \cdot T^2 \cdot \text{DNA} - k_{f,r} \cdot R \cdot \text{DNA} - k_{f,t,r} \cdot T^2 \cdot R \cdot \text{DNA} + k_{r,t} \cdot c_{d,t} + k_{r,r} \cdot c_{d,r} + k_{r,t,r} \cdot c_{d,t,r} \\
\frac{dm}{dt} &= s_m \cdot c_{d,r} - \delta_m \cdot m \\
\frac{dc_{d,t}}{dt} &= k_{f,t}^2 \cdot T^2 \cdot \text{DNA} - k_{r,t} \cdot c_{d,t} \\
\frac{dc_{d,r}}{dt} &= k_{f,r} \cdot R \cdot \text{DNA} - k_{r,r} \cdot c_{d,r} \\
\frac{dc_{d,t,r}}{dt} &= k_{f,t,r} \cdot T^2 \cdot R \cdot \text{DNA} - k_{r,t,r} \cdot c_{d,t,r} \\
\frac{dR}{dt} &= s_r \cdot m - \delta_r \cdot R.
\end{aligned} \tag{17}$$

Now, given the conservation of DNA, that is $\frac{d\text{DNA}}{dt} + \frac{dc_{d,t}}{dt} + \frac{dc_{d,r}}{dt} = 0$, and assuming the rapid kinetics, thereby invoking the quasi-steady state assumption and solving the steady state concentrations for DNA, complexes $c_{d,t}$, $c_{d,r}$ and $c_{d,t,r}$, and mRNA, it can be shown that the FadR synthesis rate we derived for the systems with Comp and NonComp are given by:

$$\begin{aligned}
\text{Comp}: \; P_R(R,T) &= \frac{a_R K_R R}{1 + K_R R + (K_T T)^2}, \\
\text{NonComp}: \; P_R(R,T) &= \frac{a_R K_R R}{(1 + K_R R + (K_T T)^2 + K_{T,R} T^2 R)} = \frac{a_R K_R R}{(1 + K_R R)(1 + (K_T T)^2)},
\end{aligned} \tag{18}$$

where $K_T^2 = \frac{k_{f,t}^2}{k_{r,t}}$ and $K_R = \frac{k_{f,r}}{k_{r,r}}$. In assuming that the binding of FadR and TetR do not affect each other in the circuit with NonComp, then we assume that $K_{T,R} = K_T^2 \cdot K_R$, which allowed us to simplify the rate equation for NonComp as shown in Supplementary Equation (18).

#### S7.2 Determining parameters for FadR positive autoregulation

For the model parameters, a lot were taken from Supplementary Table 2. However, in the spirit of using parameter values of characterised parts from the literature, we determined the parameter value representing the affinity of FadR to bind the operator on its own promoter and activate its expression, i.e. strength of positive autoregulation ($K_R$) in Supplementary Equation (18), using the dose-response data characterising the FadR-activated $P_{\text{fadRpo}}$ promoter engineered in *E. coli* DH1 in [2] (Supplementary Fig. 5(a)).

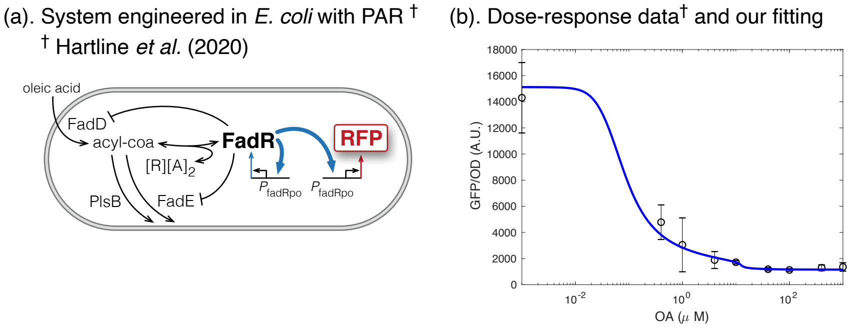

Figure 5: **Using literature data to determine parameter $K_R$ for FadR PAR Supplementary Equation (18).** (a). Schematic of the system in *E. coli* DH1 strain, from the study of [2] where FadR promoter was engineered so its expression is positively autoregulated. The engineered promoter was named $P_{\text{fadRpo}}$. (b). Plot of the average and standard deviation of the dose-response data from [2] (dots and error bars), and our optimised fitting with model Supplementary Equation (19).

The system in which the PAR was engineered is slightly different to that which we consider, i.e. our system of interest has fadE knocked out whereas the engineered system has FadR-regulated expression FadE. We therefore wrote down a dynamical model of the fadE-present system, and used it to fit the simulated dose-response to the data. We define this model as:

$$
\begin{aligned}
\frac{dR}{dt} &= b_R + \frac{a_R K_R R}{1 + K_R R} - k_f A^2 R + k_r C - \lambda R, \\
\frac{dD}{dt} &= b_D + \frac{a_D}{1 + (K_D R)^2} - \lambda D, \\
\frac{dE}{dt} &= b_E + \frac{a_E}{1 + K_E R} - \lambda E, \\
\frac{dA}{dt} &= \frac{k_{cat,D} \cdot \mathrm{OA}}{K_{m,D} + \mathrm{OA}} \cdot D - \frac{k_{cat,E} \cdot A}{K_{m,E} + A} \cdot E - \frac{k_{cat,B} \cdot A}{K_{m,B} + A} \cdot B - 2 \cdot \left(k_f A^2 R - k_r C\right) - \lambda A, \\
\frac{dC}{dt} &= k_f A^2 R - k_r C - \lambda C,
\end{aligned}
$$

where all variables are as defined in our main model, except the concentrations of FadE is defined by $E$, and growth rate is assumed constant and unaffected by the state of the system. We also model the dynamics of the RFP fluorescent reporter denoted $F$, as follows:

$$
\frac{dF}{dt} = s \cdot \left(b_R + \frac{a_R K_R R}{1 + K_R R}\right) - \lambda F,
$$

for some scaling factor $s$. It is important to note that the parameters for the expression and regulation of RFP are the same as those of FadR, as both have same promoters in lab constructs (Supplementary Fig. 5(a)).

To fit this model to the data, we first evaluated our model at steady state for various values of oleic acid, simulated the dose-response curve $F_s(\mathrm{OA})$, and performed a least-squares fitting to the data $F_d(\mathrm{OA})$ weighted by the inverse of the error of the $i^{\mathrm{th}}$ data point $\sigma_i$. That is we solved the following optimisation problem:

$$
\begin{aligned}
\min\ c \quad &= \quad \sum_i \left(\frac{F_s(\mathrm{OA}(i)) - F_d(\mathrm{OA}(i))}{\sigma_i}\right)^2, \\
\text{given constraints} \quad & \quad 0.9 \cdot \underline{p_H} \leq \underline{p} \leq 1.1 \cdot \underline{p_H}, \\
& \quad \mu_\lambda - 1.96 \cdot \sigma_\lambda \leq \lambda \leq \mu_\lambda + 1.96 \cdot \sigma_\lambda,
\end{aligned}
$$

where $\underline{p_H}$ is the vector of parameter values reported in [2], $\underline{p}$ is the vector of parameters of our model to be found, and $\mu_\lambda$ and $\sigma_\lambda$ are the mean and standard deviation of the growth rate reported in [2]. To solve this problem, we took two steps: (i) solved as a global optimization problem using genetic algorithm (`ga` in MATLAB Global optimization toolbox), and after 1000 generations, (ii) used end solution from `ga` to initialise and solve with local optimizer `fmincon`. Supplementary Table 3 summarizes the optimal parameter set found after 10 iterations of this process.

Table 3: **Optimal parameters for PAR.** From fitting model to dose-response data in Supplementary Fig. 5.

| | | | | | | |
|---|---|---|---|---|---|---|
| $\lambda$ (h$^{-1}$) | $b_R$ ($\mu$M$\cdot$ h$^{-1}$) | $a_R$ ($\mu$M$\cdot$ h$^{-1}$) | $K_R$ ($\mu$M$^{-1}$) | $b_D$ ($\mu$M$\cdot$ h$^{-1}$) | $a_D$ ($\mu$M$\cdot$ h$^{-1}$) | $K_D$ ($\mu$M$^{-1}$) |
| 0.4427 | 0.00049 | 0.1409 | 4.9115 | 0.0111 | 0.0542 | 249.7274 |
| $b_E$ ($\mu$M$\cdot$ h$^{-1}$) | $a_E$ ($\mu$M$\cdot$ h$^{-1}$) | $K_E$ ($\mu$M$^{-1}$) | $B$ ($\mu$M) | $k_{\mathrm{cat,D}}$ (h$^{-1}$) | $K_{\mathrm{m,D}}$ ($\mu$M) | $k_f$ ($\mu$M$^{-2}\cdot$h$^{-1}$) |
| 0.0046 | 47.7611 | 0.00034 | 0.1369 | 41.8052 | 0.0778 | 612.5518 |
| $k_r$ (h$^{-1}$) | $s$ | $k_{\mathrm{cat,E}}$ (h$^{-1}$) | $K_{\mathrm{m,E}}$ ($\mu$M) | $k_{\mathrm{cat,B}}$ (h$^{-1}$) | $K_{\mathrm{m,B}}$ ($\mu$M) | |
| 900.7349 | 128550 | 4.4324 | 0.102 | 194.1591 | 843790 | |

##### S7.3 Model parameters and proposal for engineering an irreversible switch

Please see Supplementary Table 4.

Table 4: **Parameters for model of irreversible switch and example engineering for its creation in lab.** Parameters values of the model of the control circuit illustrated in main text Fig. 3(a). Parameter values in the second column are based on characterised parts from the literature, with the source references in fifth column. The two last columns detail an example proposal for how to modify key system parameters our model analysis presented in the third results subsection predicts will achieve an irreversible switch.

| Characterized parts from literature | | | | | Engineering irreversibility | |
|---|---|---|---|---|---|---|
| Name | Term | Value | Units | Source | Param scaling | How |
| Growth rate | $\lambda_{\text{max}}$ | 0.1818 | $\text{h}^{-1}$ | [2], *E. coli* DH1$\Delta$*fadE* strain. | 1 | - |
| Scaled drop in growth | $s_T$ | 0.7 | - | Determined from [3], *pgi* down-regulation. | 1 | - |
| FadR leaky expression | $b_R$ | 0.0005 † | $\mu\text{M}\cdot\ \text{h}^{-1}$ | Determined in Supplementary Note S7.2, based on dose-response data of PA-FadR reporter strain from [2]. † Values are an average from 20 iterations of the fitting. | 1 | - |
| FadR promoter strength | $a_R$ | 0.1409 † | $\mu\text{M}\cdot\ \text{h}^{-1}$ | | 3.1623 | Replace RNAP binding site with T7 phage promoter $P_{\text{A1}}$, as in [4]. Then mutate to scale down to right strength. |
| FadR affinity to own promoter | $K_R$ | 4.9114 † | $\mu\text{M}^{-1}$ | | 1 | - |
| FadD leaky expression | $b_D$ | 0.0108 | $\mu\text{M}\cdot\ \text{h}^{-1}$ | [2] | 1 | - |
| FadD promoter strength | $a_D$ | 0.0517 | $\mu\text{M}\cdot\ \text{h}^{-1}$ | [2] | 1 | - |
| FadR affinity to *fadD* | $K_D$ | 305.95 | $\mu\text{M}^{-1}$ | [2] | 1 | - |
| FadD turnover rate | $k_{\text{cat,D}}$ | 49 | $\text{h}^{-1}$ | [2] | 1 | - |
| Michaelis constant | $K_{\text{m,D}}$ | 0.0672 | $\mu\text{M}$ | [2] | 1 | - |
| PlsB turnover rate | $k_{\text{cat,B}}$ | 192.91 | $\text{h}^{-1}$ | [2] | 1 | - |
| Michaelis constant | $K_{\text{m,B}}$ | 45429 | $\mu\text{M}$ | [2] | 1 | - |
| PlsB concentration | $B$ | 0.1369 | $\mu\text{M}$ | [2] | 1 | - |
| Forward sequestration | $k_f$ | 612.55 | $\mu\text{M}^{-2}\cdot\text{h}^{-1}$ | [2] | 1 | - |
| Reverse sequestration | $k_r$ | 900.73 | $\text{h}^{-1}$ | [2] | 1 | - |
| $E_g$ promoter strength | $a_g$ | $\lambda_{\text{max}}$ | $\mu\text{M}\cdot\ \text{h}^{-1}$ | This study, to ensure full expression at $\lambda_{\text{max}}$. | 1 | - |
| FadR affinity to $E_g$ promoter | $K_g$ | $= K_R =$ 4.9114 | $\mu\text{M}^{-1}$ | Assume designed with FadR operator side on *fadD* promoter; value from [2] | 1 | - |
| TetR leakiness | $b_T$ | $= b_R =$ 0.0108 | $\mu\text{M}\cdot\ \text{h}^{-1}$ | Assumed designed with *fadD* RNAP sites; value from [2]. | 1 | - |
| TetR promoter strength | $a_T$ | $= a_R =$ 0.0517 | $\mu\text{M}\cdot\ \text{h}^{-1}$ | Assumed designed with *fadD* RNAP sites; value from [2]. | 5.0119 | Replace RNAP binding sites with phage T7 promoter, $\text{P}_{\text{A1}}$ from [4], then mutate to scale down to right strength. |
| FadR affinity to *tetR* | $K_{Ri}$ | $=K_D=$ 305.95 | $\mu\text{M}^{-1}$ | Assume designed with operator site for FadR on *fadD*; value from [2]. | 0.1259 | Mutate operator to reduce affinity to right scale. |
| TetR affinity to *fadR* | $K_T$ | 5600 | $\mu\text{M}^{-1}$ | [8] | 0.0040 | Mutate operator to reduce affinity to right scale. |

#### S7.4 Determining the steady states and their stability

To determine steady states, we evaluated Supplementary Equation (14) and Supplementary Equation (1) at derivates equal to zero, and derived the nullclines:

$$N_1:\ A^2 = (b_R + P_R(R,T) - \lambda R)\cdot \frac{k_r+\lambda}{k_f R\lambda} \tag{19}$$

$$N_2:\ 0 = \frac{k_{cat,D}\cdot \mathrm{OA}}{K_{m,D}+\mathrm{OA}}\cdot D(R) - \frac{k_{cat,B}\cdot B}{K_{m,B}+A}\cdot A - 2\cdot\frac{k_f A^2 R\lambda}{k_r+\lambda} - \lambda A, \tag{20}$$

$$\text{where } D(R) = \frac{1}{\lambda}\cdot\left(b_D \frac{a_D}{1+(K_D R)^2}\right). \tag{21}$$

The steady states lie at the intersection between the two nullclines, which we determined numerically using MATLAB 2018a.

To determine the stability nature of each of the steady states, we evaluate the Jacobian matrix at the steady state and solve for the eigenvalues. If the real part of all eigenvalues is negative or zero, with at least one non-zero, then we define that steady state to be stable. If at least one eigenvalue has a non-zero positive real part, we define that steady state to be unstable. It can be shown that the Jacobian matrix of our system illustrated in main text Fig. 3(a) is:

$$J = \begin{bmatrix} \frac{\partial P_R}{\partial R} - k_f A^2 - \lambda, & 0, & -2k_f AR, & k_r, & -\lambda_{\max} s_T R & \frac{\partial P_R}{\partial T} \\ -\frac{2a_D K_D^2 R}{(1+(K_D R)^2)^2}, & -\lambda, & 0, & 0, & -D\lambda_{\max} s_T & 0 \\ -2k_f A^2, & \frac{k_{\mathrm{cat,D}}\mathrm{OA}}{K_{\mathrm{m,D}}+\mathrm{OA}}, & -\frac{k_{\mathrm{cat,B}} B K_{\mathrm{m,B}}}{(K_{\mathrm{m,B}}+A)^2} - 4k_f AR - \lambda, & 2k_r, & -A\lambda_{\max} s_T & 0 \\ k_f A^2, & 0, & 2k_f AR, & -k_r - \lambda, & -C\lambda_{\max} s_T & 0 \\ \frac{a_g K_g}{(1+K_g R)^2}, & 0, & 0, & 0, & -2\lambda_{\max} E_g s_T + \lambda_{\max}(s_T - 1) & 0 \\ -\frac{2a_T K_{Ri}^2 R}{(1+(K_{Ri} R)^2)^2}, & 0, & 0, & 0, & -\lambda_{\max} s_T T & -\lambda \end{bmatrix}, \tag{22}$$

where $\lambda$ is given by Supplementary Equation (1) and $\frac{\partial P_R}{\partial R}$ and $\frac{\partial P_R}{\partial T}$ are the partial derivates of $P_R(R,T)$ in Supplementary Equation (15) and Supplementary Equation (18).

#### S7.5 Determining which parameters affect the ability of the system to behave irreversibly

Having redesigned the control circuit, we want to understand how to tune its parameters to achieve an irreversible switch. Not all parameters may need to be modified, so we ask which parameters affect the ability of the system to behave irreversibly? Mathematically, the switch is irreversible if it can achieve at least two stable steady states when there is no inducer. To address our question we therefore evaluate the nullclines of our system, Supplementary Equation (19)–(21), at OA = A = C = 0$\mu$M. It can be shown that this gives:

$$0 = b_R + P_R(R, T(R)) - \lambda R, \tag{23}$$

where:

$$\begin{aligned} P_R &\quad \text{is as given in Supplementary Equation (15) and (18),} \\ T(R) &= \frac{1}{\lambda}\cdot\left(b_T \frac{a_T}{1+(K_{Ri}R)^2}\right), \\ \lambda &= \frac{1}{2}\cdot\left(\lambda_{\max}(1-s_T) + sqrt\lambda_{\max}^2(1-s_T)^2 + \frac{4\lambda_{\max} s_T a_g K_g R}{1+K_g R}\right). \end{aligned}$$

Though it is difficult to solve this expression analytically to find the steady states of FadR ($R$) in terms of only the system parameters, we note that their determination is not dependent on all model parameters. We infer that if more than two stable steady states can be achieved, then that depends only on engineering the system to find the correct parameter regime from amongst the following parameters $b_R, a_R, K_R, b_T, a_T, K_{Ri}, K_T$. Though there is also a dependence on parameters $a_g, K_g, \lambda_{\max}, s_T$, it is difficult to experimentally tune these. For the purposes of our analysis in the paper, we assume these latter parameters to be fixed.

### S8 Local sensitivity analysis of irreversible switch dose-response to parameters

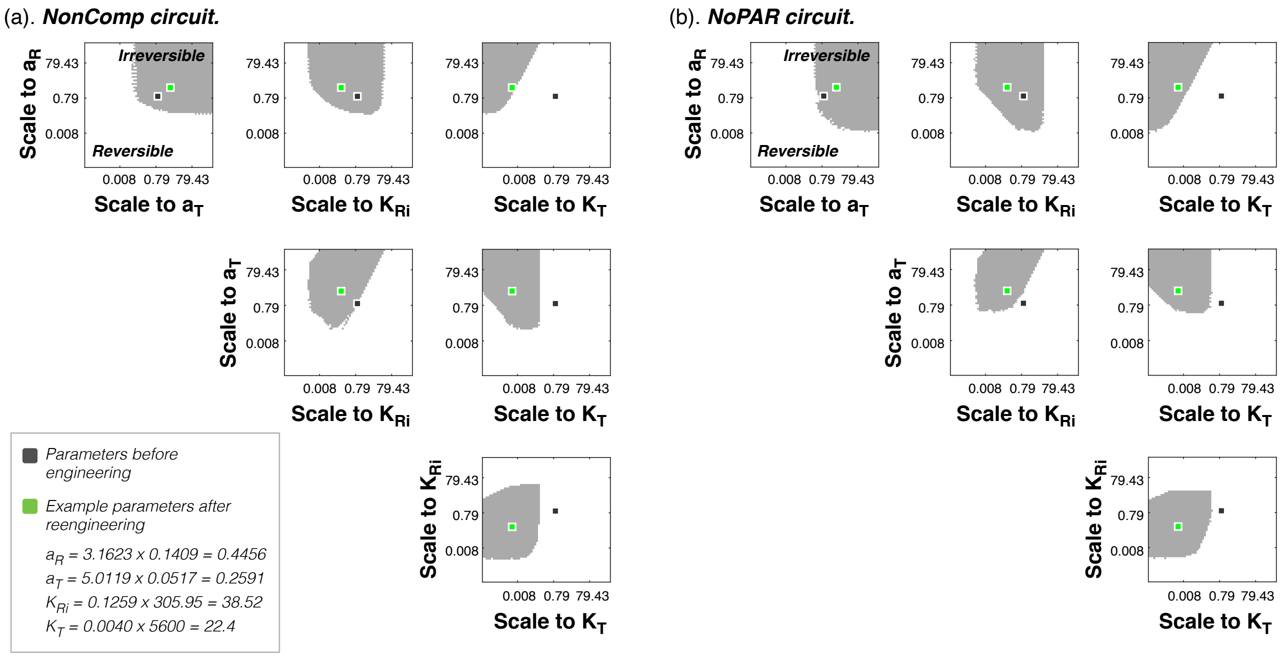

Figure 6: **Projection of the design space where the switch with given circuit topology can be irreversible,** for control circuit where FadR and TetR bind non-competitively to *fadR* promoter (a), and where FadR is not positively autoregulated (b). All parameters are tuned at the same time

### S9 How Pareto of induction regimes is affected by tuning parameters of irreversible switch

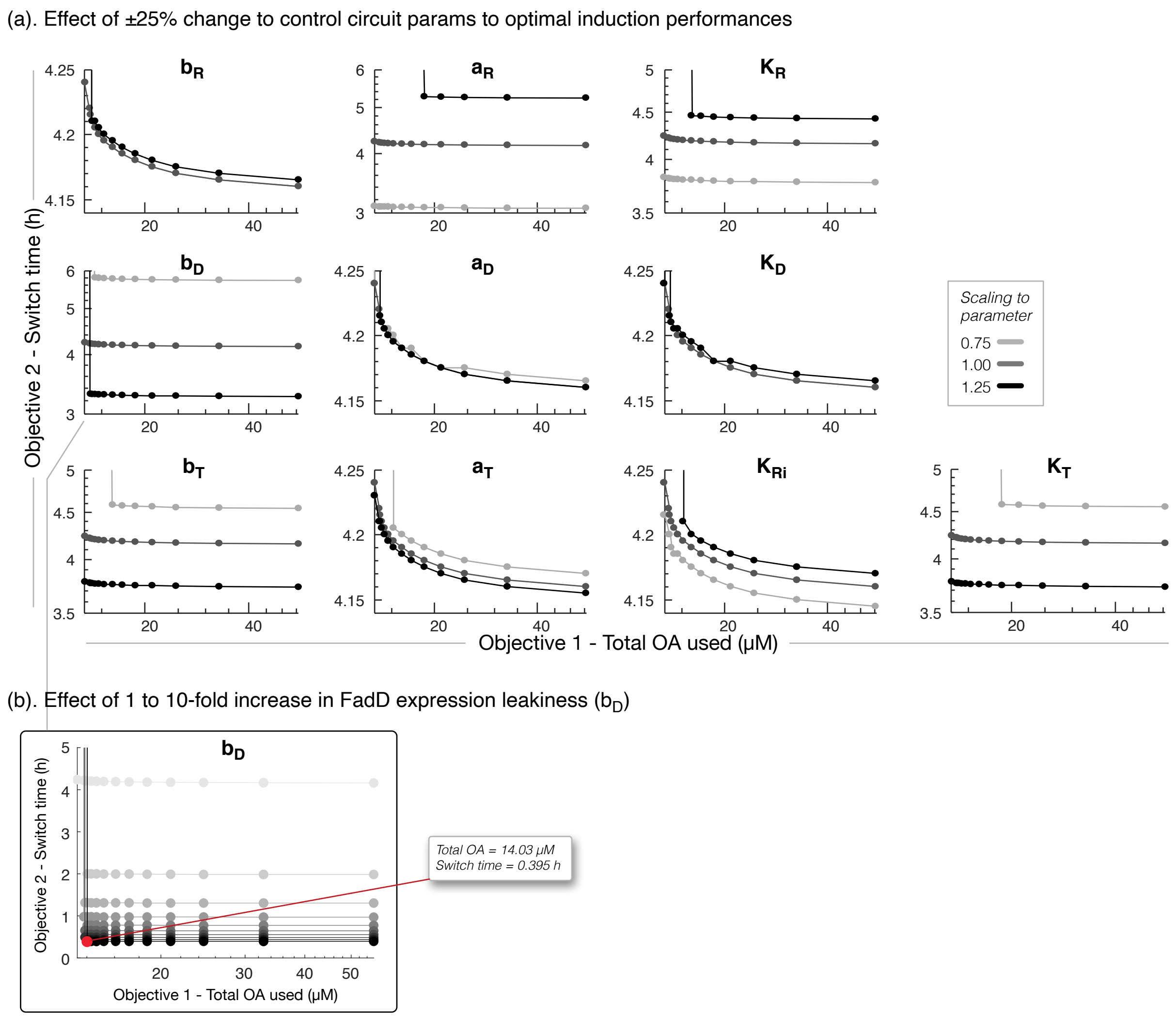

Figure 7: (a). Effect of ±25% change in experimentally-accessible parameters of our irreversible switch to the optimal induction performances found on the Pareto front of Fig. 4(b) in main text. (b). Effect of increasing FadD expression leakiness ($b_D$) from 1 to 10 times nominal value, on induction performance objectives. We indicate the optimal performance closest to the ideal point (0,0) by a red dot and report the performance objective values. There is a fundamental minimum of around 0.4 hours on the switch time achievable for increases in $b_D$.

#### S10 Performance analysis of the genetic toggle-switch and its control on growth

A commonly applied inducible switch is the genetic toggle switch [9]. Here we consider the toggle-switch, but now only induced with single inducer IPTG as opposed to two inducers IPTG and aTc. We study its application to control the switch from the growth to production phenotype, as an alternative to the proposed irreversible metabolic in this study, and present analysis of its performance. Ultimately, though we find that both switches can be engineered to behave irreversibly, and their performance is similar in terms of total inducer usage, the fact that IPTG is almost four times higher in cost outweighs the benefits of its application, relative to the oleic acid-inducible irreversible switch we presented.

##### S10.1 Model formulation

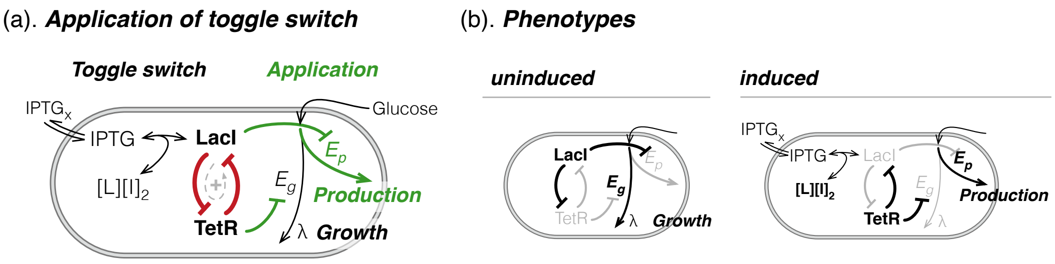

Figure 8: **Schematic of the model of the genetic toggle-switch applied as a dynamic controller.** (a). Circuitry we model to control growth and production. (b) Highlighting components in relatively high concentrations (black) when the system is uninduced and induced, exhibiting the growth (left) and production phenotypes (right).

**Equations.** We model the genetic toggle switch and its application as a dynamic controller, to switch from the growth to production phenotype for inductions of IPTG (Supplementary Fig. 8(a)), with the following set of ordinary differential equations:

$$\begin{aligned}
\frac{dI_x}{dt} &= I_{\text{feed}} - k_i(I_x - I) \\
\frac{dI}{dt} &= k_i \cdot (I_x - I) - 2 \cdot (k_f I^2 L - k_r C) - \lambda(E_g)I, \\
\frac{dL}{dt} &= b_L + \frac{a_L}{1 + (K_L T)^2} - (k_f I^2 L - k_r C) - \lambda(E_g)L, \\
\frac{dC}{dt} &= (k_f I^2 L - k_r C) - \lambda(E_g)C, \\
\frac{dT}{dt} &= b_T + \frac{a_T}{1 + K_T L} - \lambda(E_g)T, \\
\frac{dE_g}{dt} &= \frac{a_g}{1 + (K_g T)^2} - \lambda(E_g) \cdot E_g,
\end{aligned} \tag{24}$$

where $\lambda(E_g) = \lambda_{\max} \cdot (E_g s_T - s_T + 1)$, as in Supplementary Equation (1), and $I_x, I, L, C, T, E_g$ are concentrations, in units of $\mu$M, of IPTG in media, IPTG in a cell, LacI, sequestered complex of a LacI dimer to two IPTGs, TetR, and the growth associated enzyme, respectively. To capture the cooperativity effect of LacI repression [9], we fixed Hill coefficient of 2 in the third and sixth equations (repression of LacI and $E_g$ expressions).

**Parameters.** Please see Supplementary Table 5 for the full list of parameters, their values and units. To enable a fair comparison between the application of the toggle-switch and the irreversible switch in Supplementary Note S7, we set parameters of the expression of LacI and TetR similar to those defined for the

Table 5: **Parameters of the model in Supplementary Fig. 8(a) of the toggle-switch applied for dynamic control of growth and production.**

| Name | Term | Value | Units | Source |
|---|---|---|---|---|
| Growth rate | $\lambda_{\max}$ | 0.1818 | $h^{-1}$ | same as Supplementary Table 4. |
| Scaled drop in growth | $s_T$ | 0.7 | - | same as Supplementary Table 4. |
| IPTG feed in flux | $I_{\text{feed}}$ | - | $\mu M \cdot h^{-1}$ | to optimise |
| IPTG feed in time | $T_{\text{exp}}$ | - | h | to optimise |
| free diffusion rate | $k_i$ | 3600 | $h^{-1}$ | set in this study. |
| Sequestration fwd rate | $k_f$ | 1057.7 | $\mu M^{-2} \cdot h^{-1}$ | fitting, Supplementary Fig. 9(b) |
| Sequestration rev rate | $k_r$ | 1292.1 | $h^{-1}$ | fitting, Supplementary Fig. 9(b) |
| LacI leaky expression | $b_L$ | 0.0005 | $\mu M \cdot h^{-1}$ | $= b_R$, Supplementary Table 4. |
| LacI promoter strength | $a_L$ | 0.5347 | $\mu M \cdot h^{-1}$ | = 0.14 * 3.16 * 1.2, $= 1.2 * a_R$ in Supplementary Table 4. |
| Affinity of TetR to inhibit LacI exp | $K_L$ | 100 | $\mu M^{-1}$ | set in this study. |
| TetR leaky expression | $b_T$ | 0.0011 | $\mu M \cdot h^{-1}$ | $= 0.1 * b_T$ in Supplementary Table 4. |
| TetR promoter strength | $a_T$ | 0.2591 | $\mu M \cdot h^{-1}$ | = 0.05 * 5.01 $= a_T$, Supplementary Table 4. |
| Affinity of LacI to inhibit TetR exp | $K_T$ | 305.95 | $\mu M^{-1}$ | $= K_{Ri}$, Supplementary Table 4. |
| $E_g$ promoter strength | $a_g$ | $\lambda_{\max}$ | $\mu M \cdot h^{-1}$ | same as Supplementary Table 4. |
| Affinity of TetR to inhibit $E_g$ exp | $K_g$ | 100 | $\mu M^{-1}$ | $= K_L$, set in this study. |

irreversible switch (see Supplementary Table 4) to achieve similar steady state concentration levels (i.e. similar expression burden).

We needed to determine the specific kinetics of the reversible binding and sequestration of LacI by IPTG. It is important to note that we model the expression of LacI as a dimer, and assume that each LacI dimer binds to a single synthetic operator site, and that two molecules of IPTG bind to sequester each LacI dimer, emulating the experimental setup in [4]. Supplementary Fig. 9(a) shows the model we used to capture the dynamics of LacI-IPTG sequestration, and we employed least-squares fitting (using `fmincon` from the MATLAB 2018a Optimization Suite) to determine the optimal values of the forward and reverse binding rates $k_f$ and $k_r$, that minimises the Euclidean distance between simulations of the model in (a) (bold line, Supplementary Fig. 9(b)) and experimental data taken from [10]. Please see Supplementary Fig. 9 caption for details of experimental data used.

#### S10.2 Finding parameter space where toggle switch seems to behave irreversibly

One can modify the toggle-switch behaviour in the lab by modifying promoter strengths of LacI and TetR ($a_L$ and $a_T$) via mutating their RNAP binding sequences, and the affinity they bind to their respective operators to repress the others expression ($K_L$ and $K_T$) via mutating the operator sequences. The study of [9] reported that for some modifications, the toggle-switch was observed to behave similarly to an irreversible switch, i.e. the induced state remained stable after a temporal IPTG induction and its washout.To uncover the parameter space were this is possible we explored the $a_L, a_T, K_L, K_T$-parameter space by scaling their nominal values in Supplementary Table 5 by $10^{-3}$ to $10^3$.

In fact, the dose-response curve of free LacI concentration vs titrations of IPTG inductions based on the nominal parameter values already showed an effectively irreversible toggle-switch (Supplementary Fig. 10(a)). Projections of the 4D-space of the four parameters explored (Supplementary Fig. 10(b)) unveil similar principles of how to engineer the toggle-switch to become irreversible, as those found for the oleic acid-inducible switch, i.e. stronger promoter strengths but weaker affinities of each TF to inhibit the others expression.

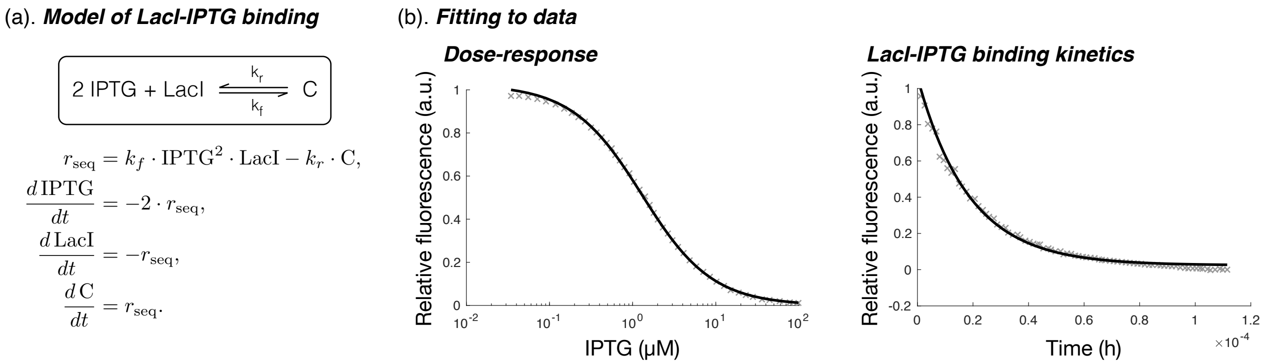

Figure 9: **Determining parameters of IPTG sequestration of LacI.** (a). Chemical and mathematical equations of LacI sequestration by gratuitous inducer IPTG, assuming 2 molecules of IPTG bind to each dimer of LacI. (b). Least-squares fit of model to two data sets (gray crosses) from Xu and Matthews [10], with optimal values of forward and reverse rates of sequestration found as $k_f = 1057.7\ 1/(\mu\text{M}^2{\cdot}\text{h})$ and $k_r = 1292.1\ \text{h}^{-1}$. The left plot shows data are of the steady state relative fluorescence measured for titrations of IPTG added to $1.5 \times 10^{-7}$M LacI monomer. The right plot shows data of the decrease in relative fluorescence measured by rapid stop flow method after addition of 0.05mM IPTG to $1 \times 10^{-6}$M LacI monomer. Important note: it is assumed that relative fluorescence is representative of the number of free LacI in solution.

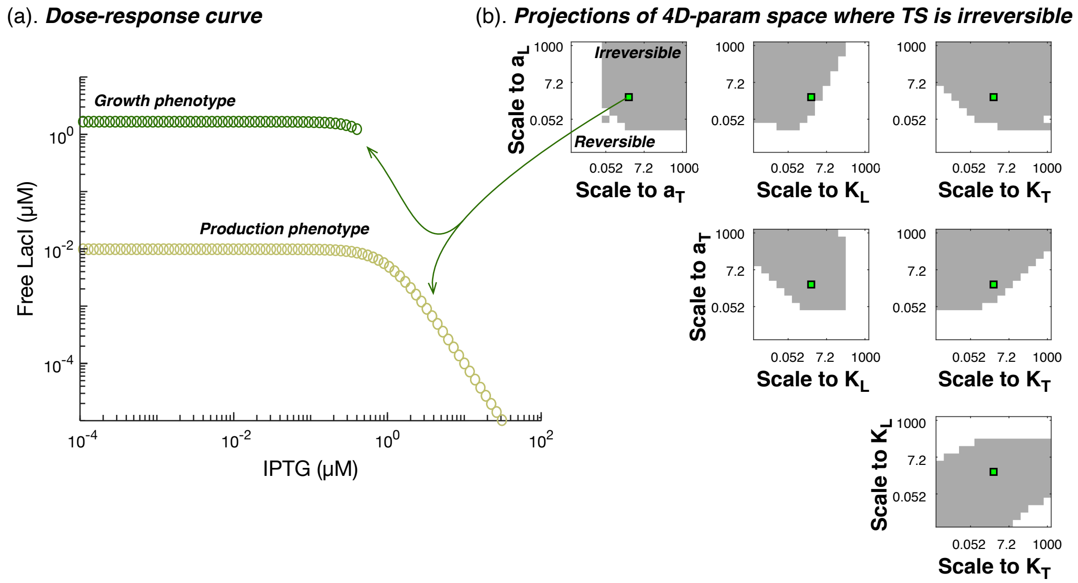

Figure 10: **Exploring param space unveils design principles for engineering irreversible toggle-switch** (a). Dose-response curve of the concentration of free LacI for titrations of IPTG, for the nominal parameter set in Supplementary Table 5. (b). Projections of 4D-parameter space onto planes of two parameter pairs, highlighting parameter regimes where the toggle-switch can behave irreversibly.

#### S10.3 Optimizing induction regime to minimise inducer use and switch time

Using the parameters that define the irreversible toggle-switch (Supplementary Table 5), we searched for the optimal induction regime that would minimise the total use of IPTG and switch time, as was sought for the oleic acid-inducible irreversible switch. That is, we are again solving the multiobjective optimisation defined as:

$$
\begin{aligned}
&\min_{I_{\text{feed}},T_{\text{exp}}} \ (J_1, J_2)\,,\\
&J_1 = \int_{t=0}^{100} f_{\text{in}}(t)\,\mathrm{d}t; \quad J_2 = \min\left(t \mid \text{LacI}(t) \le 0.05\mu M\right);\\
&\text{where } f_{\text{in}}(t) = \begin{cases} I_{\text{feed}}, & \text{for } t \le T_{\text{exp}},\\ 0, & \text{for } t > T_{\text{exp}},\end{cases}\\
&\text{and subject to} \begin{cases} 1\mu\text{M}\cdot\text{h}^{-1} \le I_{\text{feed}} \le 50\mu\text{M}\cdot\text{h}^{-1},\\ 0.1\text{h} \le T_{\text{exp}} \le 5\text{h},\end{cases}
\end{aligned}
\tag{25}
$$

where we search over the two induction regime *tuning dials*: flux of IPTG feed-in $I_{\text{feed}}$, and time length of feed in (also referred to as the exposure time) $T_{\text{exp}}$.

In solving Supplementary Equation (4) (using MATLAB 2018a Global Optimization suite solver `gamultiobj`, see Methods in main text), we obtained a Pareto front of optimal solutions, as shown in Supplementary Fig. 11(a, left plot), indicative of the trade-off between the two objectives. We find that total IPTG used is of a similar order of magnitude as total oleic acid used in the oleic acid-inducible switch (Supplementary Fig. 11(a)-left vs Fig. 4(b) in main text). However, interestingly, though reducing primarily IPTG feed in flux by just less that 40% (without significantly changing exposure time) (Supplementary Fig. 11(a), right) can reduce total IPTG usage by around 25%, it comes at a substantial cost of increased switch times, by over 300-fold. Looking at the induction dynamics based on the Pareto solution lying closest to the ideal point

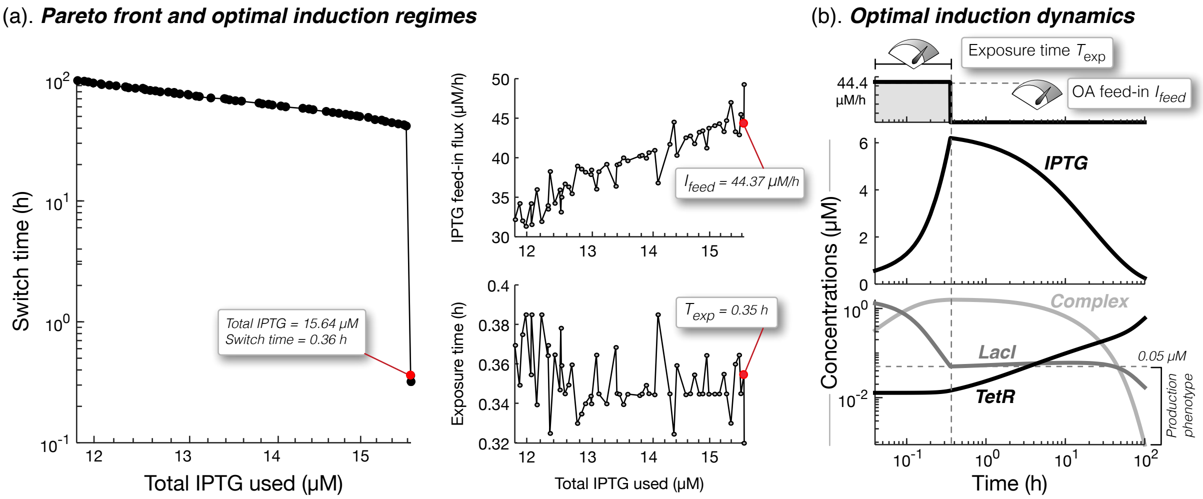

Figure 11: **Pareto solutions of optimal induction regimes to minimise total IPTG usage and switch times.** (a). Pareto front of objective values found (left plot), with plots of the IPTG feed in flux and exposure times corresponding to each Pareto solution (right plots). The solution lying closest to the ideal point (0,0) is highlighted as a red dot, and the respective optimal objective values and induction parameter values are shown in boxes. (b). Time course simulations using the optimal induction regime lying closest to the ideal point.

(Supplementary Fig. 11, red dots), we observe the expected accumulation of IPTG during inducer exposure, but interestingly, it is very slowly diluted away by a non-zero growth rate of the cell (growth reduced to 0.055 $\text{h}^{-1}$). It is also interesting to note that after induction ceases, free LacI concentrations slightly rise before finally falling due to a combination of increased expression of its repressor TetR and dilution by cell growth.

#### S10.4 Effect of change in circuit parameters on Pareto front

To understand how the Pareto front is changed for modifications in the circuit parameters, we recomputed both performance objective values for all solutions along the Pareto front for ±10% change in the following respective circuit parameters: $b_L, a_L, K_L, b_T, a_T, K_T$.

As shown in Supplementary Fig. 12(a), if we decrease LacI concentration, by decreasing promoter strength ($a_L$) or increasing inhibition of its expression by TetR ($K_L$), or increase TetR concentration, by increasing its expression ($a_T$ or $b_T$) or decreasing inhibition of its expression by LacI ($K_T$), then we can drastically decrease the switch time for many of the Pareto induction regimes. Nevertheless, since this objective is significantly outperformed by the oleic acid-inducible switch, we find that the advantage of tuning the toggle switch is being able to reduce total IPTG usage. However, even if we decrease LacI expression ($a_L$) to 10% or increase TetR expression ($a_T$) by 10 times (Supplementary Fig. 12(b)), there is a fundamental limit to the minimum total IPTG usage, which is ≈12$\mu$M.

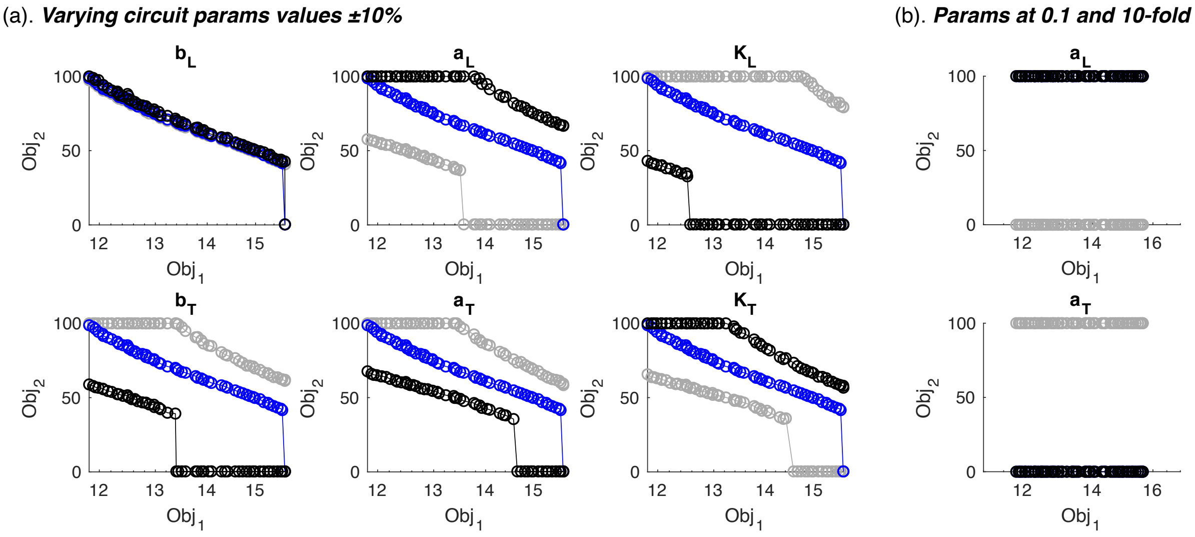

Figure 12: **Change in Pareto solutions.** (a). For ±10% change in respective value of experimentally-accessible circuit parameters. Data in gray, blue and black represent the recalculated objectives after scaling the respective nominal parameter values shown in Table Supplementary Table 5 by 0.9, 1 and 1.1, respectively. (b) For scaling promoter strengths from 10% to 10 times their nominal values shows limit on minimum achievable total IPTG use (Obj1) of 11.8$\mu$M.

[10] Xu, J. & Matthews, K. S. Flexibility in the Inducer Binding Region is Crucial for Allostery in the Escherichia coli Lactose Repressor. *Bone* **48**, 4988–4998. `NIHMS150003`.
